# Combined Vapor Exposure to THC and Alcohol in Pregnant Rats: Maternal Outcomes and Pharmacokinetic Effects

**DOI:** 10.1101/2020.07.24.220103

**Authors:** Kristen R. Breit, Cristina Rodriguez, Annie Lei, Jennifer D. Thomas

**Affiliations:** Center for Behavioral Teratology, Dept. of Psychology, San Diego State University

**Keywords:** alcohol, THC, vapor, prenatal, maternal, pharmacokinetics

## Abstract

Cannabis is the most frequently used illicit drug among pregnant women, yet the potential consequences of prenatal cannabis exposure on development are not well understood. Electronic cigarettes have become an increasingly popular route of administration among pregnant women, in part to user’s perception that e-cigarettes are a safer route for consuming cannabis products. Importantly, half of pregnant women who consume cannabis also report consuming alcohol, but research investigating co-consumption of these drugs is limited, particularly with current routes of administration. The purpose of this study was to establish a co-exposure vapor inhalation model of alcohol and THC in pregnant rats, to ultimately determine the effects on fetal development. Pregnant Sprague-Dawley rats were exposed to moderate doses of THC via e-cigarettes, alcohol, the combination, or vehicle daily from gestational days 5-20. Importantly, pharmacokinetic interactions of alcohol and THC were observed during pregnancy. Combined exposure consistently increased blood alcohol concentrations, indicating that THC alters alcohol metabolism. In addition, THC levels also increased over the course of pregnancy and THC metabolism was altered by alcohol. Alcohol, but not THC, exposure during pregnancy reduced maternal weight gain, despite no group differences in food intake. Neither prenatal alcohol nor THC exposure altered gestational length, litter size, sex ratio or birth weight. However, prenatal alcohol exposure delayed eye opening, and prenatal THC exposure decreased body weights during adolescence among offspring. These individual and synergistic effects suggest that this novel co-exposure vapor inhalation paradigm can effectively be used to expose pregnant dams, exerting some effects on fetal development, while avoiding nutritional confounds, birth complications, or changes in litter size. With this model, we have demonstrated that combining THC and alcohol alters drug metabolism, which could have important consequences on prenatal development.

## 1 Introduction

Prenatal alcohol exposure can disrupt physical, neurological, and behavioral development, leading to a range of outcomes known as fetal alcohol spectrum disorders (FASD). Individuals with FASD may exhibit impairments in a number of behavioral/cognitive domains, including learning, attention, executive functioning, emotional regulation, social interactions, motor coordination, and impulse control, which can lead to serious problems in school and daily life (Khoury et al., 2015; Norman et al., 2013). Prenatal alcohol exposure can also induce growth deficits and facial dysmorphology; individuals with neural impairment, poor growth and craniofacial abnormalities may be diagnosed with fetal alcohol syndrome, which lies on the most severe end of the spectrum (Jones et al., 1973). FASD pose a global health concern, as approximately 1 in 10 women report some alcohol consumption during pregnancy (Roozen et al., 2016) and prevalence rates of FASD in the U.S. range from 1-5% (May et al., 2020).

However, women may consume other drugs besides alcohol during pregnancy. Given the recent policy changes and decriminalization of cannabis products in many U.S. states and Canada, with concomitant increases in availability, cannabis use has been rising dramatically over the last decade (SAMHSA, 2020). This is of particular concern among adolescents and young adults, including women of child-bearing age (Brown et al., 2017). For instance, 20% of U.S. women ages 18-25 use cannabis (SAMHSA, 2020), with higher rates in areas where cannabis is legal (Lee et al., 2020; Reimann et al., 2011). In particular, prevalence of cannabis use among pregnant women ranges from 3-10% (Coleman-Cowger et al., 2017; Ko et al., 2015; Obisesan et al., 2020; Oh et al., 2017; SAMHSA, 2020; Volkow et al., 2019; Young-Wolff et al., 2019), with even higher rates among pregnant teens (Gupta et al., 2016).

Unfortunately, young people increasingly view all cannabis use as safe (Brown et al., 2017; Johnston et al., 2015), and both pregnant and non-pregnant women perceive cannabis as posing a minimal risk (Jarlenski et al., 2017). In fact, many pregnant women purposefully take cannabis products for pregnancy-related illness such as nausea (Dickson et al., 2018), even though cannabis may actually provoke recurrent nausea and vomiting rather than combat it (Kim et al., 2018). Cannabis consumption during pregnancy is especially prevalent during the first and second trimesters; in the U.S. 8% of pregnant women in the first trimester report consuming cannabis in the past 30 days (Volkow et al., 2019). Given that 40% of pregnancies are unplanned (Sedgh et al., 2014), even if risks were well recognized, fetal exposure to cannabis remains a serious public health concern.

However, the risks of prenatal cannabis exposure are still not well understood, despite the high prevalence of use. The results from existing prospective and retrospective clinical studies examining prenatal cannabis exposure are mixed, likely due to differences in cannabis exposure levels, prospective versus retrospective approaches, confounds of other drug use, age and nature of outcome measures, and a host of other factors. Thus far, it appears as though prenatal cannabis exposure generally does not produce physical birth defects, although it may reduce birth weight (Day et al., 1991a; Day et al., 1991b; Fergusson et al., 2002; Fried et al., 1987; Huizink, 2014; Hurd et al., 2005; Paul et al., 2019). Other clinical evidence suggests that children exposed to prenatal cannabis have altered emotional, behavioral, and cognitive development, particularly in executive functioning (Huizink, 2014). More recent clinical data from the longitudinal Adolescent Brain Cognitive Development study suggest that prenatal cannabis increases the risk for low weight at birth, as well as psychopathology symptoms and cognitive deficits among children 9-11 years of age (Paul et al., 2019).

Notably, the elucidation of the consequences of prenatal cannabis is particularly challenging given that accessibility and potency levels continue to rapidly change. For example, potency through selective cultivation of the primary psychoactive constituent of cannabis, delta-9-tetrahydrocannabinol (THC), has increased from 3.4% in 1993 to 55.7% or higher in 2017 (Chandra et al., 2019; Mehmedic et al., 2010), with the average potency being approximately 17.1% (Chandra et al., 2019). Furthermore, levels are even higher among synthetic cannabinoid preparations (e.g. Spice), compared to cultivated marijuana (Botticelli, 2016; Subbanna et al., 2013). Thus, results from current longitudinal clinical studies may not represent current cannabis consumption levels (Gunn et al., 2016; Huizink, 2014; Jaddoe et al., 2012).

In addition to changes in accessibility and potency levels of cannabis, administration routes have also evolved. Prevalence rates of general electronic cigarette (e-cigarette) use among pregnant women in the U.S. is estimated to be between 5-14% (Cardenas et al., 2019). This includes the use of e-cigarettes to consume cannabis and its constituents, as vaping has become one of the most popular routes of administration for cannabis. In fact, pregnant women, particularly young women, have the perception that consumption of cannabis through e-cigarettes (vaping) is safe, despite the recent increase of deaths in the U.S. due to Vaping-Associated Pulmonary Illness (Carlos et al., 2019) and the known dangers of traditional smoking during pregnancy (Mark et al., 2015). One recent survey illustrated that among pregnant women with equivalent knowledge about the dangers of traditional smoking during pregnancy (such as smoking cigarettes or blunts), 43% believed that vaping is a safer alternative (Mark et al., 2015). Moreover, e-cigarette use during pregnancy is likely to occur among women diagnosed with substance abuse disorders and/or among women who are trying to transfer from smoking traditional cigarettes or blunts once pregnancy is confirmed (Oncken et al., 2017). Importantly, recent studies examining various routes of administration suggest that vaping cannabis may have more detrimental effects on a developing fetus compared to other traditional routes, as higher THC concentrations can be reached both in the vaping liquid and in the consumer’s blood levels (Young-Wolff et al., 2020). Yet, little research has examined the health consequences resulting from e-cigarette use during pregnancy, despite requests from medical professionals (Brandon et al., 2015; Suter et al., 2015).

Another challenge in understanding the potential effects of cannabis exposure on fetal development is the high rate of polydrug consumption. According to the National Household Survey on Drug Abuse, half of pregnant women who report consuming cannabis also report drinking alcohol (Substance Abuse and Mental Health Services Administration, 2015). However, accurate data reflecting concurrent use of alcohol and cannabis, as well as exposure levels, among pregnant women are difficult to obtain as women who consume either alcohol or cannabis frequently under-report their usage due to stigma (Lange et al., 2014; Young-Wolff et al., 2017).

Despite the known rates of co-use, it is still relatively unclear whether THC exposure could exacerbate alcohol’s teratogenic effects, since results from both clinical and preclinical studies have been mixed (Abel et al., 1990; Goldschmidt et al., 2004). Most clinical studies focus on the effects of each drug on fetal development separately (Day et al., 1993; Fried et al., 1992; Fried et al., 1990; Richardson et al., 2002), rather than the combination of effects (Goldschmidt et al., 2004). Similarly, controlled animal models have also primarily focused only on the teratogenic effects of each drug individually, although a few have examined combined effects on physical development and/or neurotoxicity (Abel et al., 1987; Fish et al., 2019; Hansen et al., 2008).

Studies using rodent models can help provide quick, accessible information regarding the potential consequences of prenatal alcohol and THC exposure at the levels and routes currently being consumed. This study used a clinically relevant and effective rodent co-exposure model of e-cigarette THC exposure and alcohol to examine the consequences of prenatal co-exposure. Vapor inhalation paradigms that deliver THC using commercially available e-cigarettes have recently been used in rodents to expose non-pregnant rats (Javadi-Paydar et al., 2019; Javadi-Paydar et al., 2018; Nguyen et al., 2019), but the use of such a model in pregnant rats is more limited (Weimar et al., 2020). To date, a rodent vapor inhalation paradigm for co-exposure to alcohol and cannabis during pregnancy has not been utilized.

The purpose of this study was to establish a prenatal rodent vapor inhalation procedure that would 1) model maternal intake routes and levels, 2) determine the effects on physical and behavioral development of offspring and 3) determine how the effects of the combination of drugs compare to either drug alone. Using vapor inhalation and commercially available e-cigarette tanks, pregnant Sprague-Dawley rats were exposed to alcohol, THC, the combination, or a vehicle during the first and second trimester human equivalent. We hypothesized that prenatal exposure to either drug via vapor inhalation would negatively impact developmental outcomes in the offspring and that combined exposure would exacerbate these effects. Behavioral data from the offspring are reported in a separate article. This report includes the pharmacokinetic effects of co-exposure, as well as maternal effects and physical effects on the offspring.

## 2 Methods

### 2.1 Subjects

This study used a model of combined prenatal exposure to vaporized alcohol and THC via electronic cigarettes to determine the consequences of prenatal THC exposure alone and with alcohol (Figure 1). All procedures included in this study were approved by the San Diego State University (SDSU) Institutional Animal Care and Use Committee (IACUC) and are in accordance with the National Institute of Health’s *Guide for the Care and Use of Laboratory Animals*. Naïve female Sprague-Dawley rats were obtained from Charles River Laboratories (Hollister, CA) on postnatal day (PD) 60 and allowed to acclimate for at least two weeks prior to any handling or procedures in the animal care facilities at the Center for Behavioral Teratology (CBT) at SDSU.

**Figure 1.**
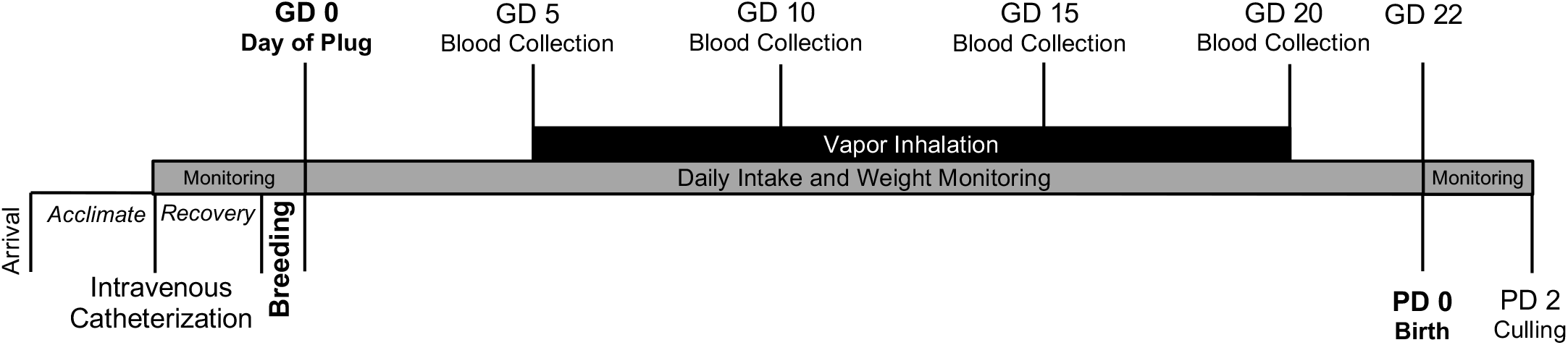
Timeline of study procedures.

#### 2.1.1 Intravenous Catheterization

Following the 2-week acclimation period, all dams were surgically implanted with an intravenous catheter in the right jugular vein to facilitate blood sampling with minimal stress throughout the vapor inhalation period. Dams were anesthetized with 4% isoflurane and had their right upper chest region and lower back area shaved. Shaved areas were sterilized with ethanol and betadine. A straight incision was made across the lower back and a small incision was made above the right jugular vein. The catheter was secured under the skin on the back, and the tubing was thread subdermally over the right shoulder and implanted into the right jugular vein. Tubing was flushed with heparinized bacteriostatic saline via syringe to ensure proper placement and then cuts were sealed with veterinary glue (VetBond; 3M). Catheters were then covered with a plastic hood and a metal screw cap to prevent chewing. Dams were administered an antibiotic (Cefazolin, 100 mg/mL; Victor Medical) and a painkiller (Flunixin, 2.5 mg/mL; Bimeda) following surgery and for 2 consecutive days post-surgery (subcutaneous injection, 0.001 mL/g). Catheterized dams were then singly housed to recover for at least 1 week prior to breeding and were monitored daily for weight loss and healing complications.

#### 2.1.2 Breeding

Following recovery from the catheter implantation, dams were paired with one male from the CBT breeding colony. Breeding pairs were housed in a standard Allentown rat cage with a raised grid wire floor; a filter paper was placed under the wire floor to catch any seminal plugs. Pairs were checked daily each morning for the presence of a seminal plug, which was designated as gestational day (GD) 0. Throughout breeding, pairs had *ad libitum* access to food and water.

### 2.2 Prenatal Vapor Inhalation Exposure

Pregnant dams were assigned to 1 of 4 prenatal exposure groups on GD 0. Pregnant dams were exposed to either vaporized ethanol (EtOH) or Air, and half of each group was either exposed to THC or the Vehicle via electronic cigarette (e-cigarette). Thus, this study used a 2 (EtOH, Air) × 2 (THC, Vehicle) design.

#### 2.2.1 Daily Monitoring

Beginning on GD 0 and throughout gestation (GD 0-22), body weights were recorded each morning before any procedures began. All dams were given free access to 200 g of standard pellet lab chow (LabDiet 5001) and 400 mL of water in a graduated bottle each day; food and water intake were recorded each morning and refilled.

#### 2.2.2 Drugs

THC for the e-cigarette tanks was obtained through the National Institutes of Drug Abuse (NIDA) Drug Supply Program and arrived in 95% ethanol. To remove the alcohol, a speedvac concentrator (Thermo Scientific, Savant SPD111V/RVT400) was used to evaporate the alcohol and the remaining THC was then dissolved in propylene glycol to achieve the desired concentration (100 mg/mL). EtOH for prenatal vaporizing was obtained from Sigma-Aldrich.

#### 2.2.3 Vapor Inhalation Paradigm

Beginning on GD 5, dams were exposed to the vapor inhalation paradigm once per day until GD 20 (equivalent to the first and second trimesters in pregnant humans). Vaporized drug exposure was administered using two 4-chamber vapor inhalation apparatuses designed by La Jolla Alcohol Research Inc (La Jolla, San Diego, CA). The vapor inhalation system uses a sealed cage identical in structure and dimensions as standard rat Allentown cages (Allentown, PA), a metered pump and heated flask to vaporize EtOH, a programmable computer-controllable adapter to trigger commercially available e-cigarette tanks (SMOK V8 X-Baby Q2), and vacuum regulation of the air/vapor flow. Each chamber contains individual airflow meters to ensure that the proper airflow is maintained consistently; all dams, drug containers, and airflow meters were monitored throughout the vapor exposure period.

Pregnant dams were exposed to either vaporized EtOH (95% at 10 L/min airflow) or Air in a constant stream for 3 hours. At the completion of the 3 hours, half of the EtOH-exposed dams were exposed to either THC (100 mg/mL at 2 L/min airflow) or the Vehicle (propylene glycol; Sigma-Aldrich) for 30 min via an e-cigarette tank. E-cigarette puffs were administered in a 6-sec puff every 5 min for 30 min (7 puffs total). All subjects were then given 10 min of additional air flow (2 L/min) to clear out any residual drug before removal from the chambers.

#### 2.2.4 Core Body Temperatures

Before and after each vapor inhalation session, core body temperatures were taken from each dam via a rectal thermometer. Previous research suggests that vaporized THC exposure via e-cigarette may reduce core body temperatures during exposure in both male and female rats (Javadi-Paydar et al., 2019; Javadi-Paydar et al., 2018; Nguyen et al., 2016).

#### 2.2.5 Blood Levels

Blood samples were taken throughout gestation to confirm drug and metabolite levels. Blood (300 µL) was collected from each dam via the intravenous catheter on GD 5, 10, 15, and 20 at 15, 30, 60, 90, and 180 minutes post-vapor inhalation. The catheter was flushed with heparinized saline before and after each collection (0.2 mL). In the rare case that blood could not be successfully drawn via the catheter, blood was instead taken via tail vein cut. Blood samples were immediately placed in the centrifuge to separate plasma and stored at −80°C until analyses were conducted.

Plasma levels (10 µL) of blood alcohol concentrations (BAC) were analyzed at the CBT using an Analox Alcohol Analyzer (4Model AMI; Analox Instruments; Lunenburg, MA). THC and metabolite levels were analyzed by MZ Biolabs (Tucson, AZ). Fifty µl of each plasma sample was precipitated (vigorous vortex and 30 min incubation at 4°C) using 200 µl HPLC grade acetonitrile containing 10 ng/mL 11-nor-9-Carboxy-Δ^9^-THC-D_9_ as an internal standard (Cerilliant T-007-1ML). The supernatant was centrifuged for 10 min (4°C) and transferred to a 96-well plate for analysis using LC-MS2/MS3. An 8-point standard curve containing Δ^9^-THC (Cerilliant T-005-1ML), 11-Hydroxy-Δ^9^-THC (Cerilliant H-026-1mL), and 11-nor-9-Carboxy-Δ^9^-THC (Cerilliant T-018-1ML) was prepared with concentrations of 3 analytes ranging from 781 pg/ml to 100 ng/mL. A Surveyor HPLC (Thermo Scientific) connected to a LTQ Velos Pro mass spectrometer (Thermo Scientific) was used to separate and quantify Δ^9^-THC and metabolites (compartment temperature of 6°C and column temperature of 25°C). A reverse phase gradient was used for elution from a 2mm × 150 mm C18 column (Phenomenex 00F-4435-B0) at a flow of 150 µl/min. Gradient conditions were 75% A, 25% B ramping to 100% B in 11 minutes, held at 100% B for 4 minutes, followed by equilibration at 75% A, 25%B for 6 minutes, where A is water containing 10 mM ammonium bicarbonate adjusted to pH 6 with formic acid and B is methanol. Eluate was analyzed by the LTQ Velos Pro using negative ions. Quantitation was performed using the Quan Browser software from Thermo Scientific. An in-house QC was prepared with 10 ng/mL each of Δ^9^-THC, 11-Hydroxy-Δ^9^-THC, and 11-nor-9-Carboxy-Δ^9^-THC prepared in drug free canine plasma. The linear quantitative range of the assay was 781 pg/ml to 100 ng/mL.

#### 2.2.6 Birth and Litter Monitoring

The day of birth (usually GD 22) was designated as PD 0 and the gestation length was recorded. On PD 2, the number of pups, pup weights, and pup sexes were recorded. Litters were culled to 8 pups (4 males and 4 females when possible). Offspring were monitored until PD 30 to record body weights and the first day of eye opening (the first day both eyes were fully open).

### 2.3 Statistical Analyses

All data were analyzed using the Statistical Packages for Social Sciences (SPSS, version 26; IBM). Data were analyzed using a 2 (EtOH: EtOH, Air) × 2 (THC: THC, Vehicle) Analysis of Variance (ANOVA) with significance values set at *p* < 0.05. Data analyzed across multiple days and/or times used a Repeated Measures ANOVA with Day and Time as within-subjects variables. Offspring data additionally used sex (female, male) as a between-subjects factor. Student Newman Keuls (SNK) post hoc tests (*p* < 0.05) were used when needed. For eye opening, nonparametric analyses were used.

## 3 Results

### 3.1 Subjects

Data analyses were conducted using only dams that completed the full vapor inhalation procedure and successfully gave birth, with final n’s of 10-13 in each exposure group (EtOH+THC: 10; EtOH+Vehicle: 12; Air+THC: 13; Air+Vehicle: 12). Twelve additional dams were excluded due to miscarriage, dystocia, or cannibalization (EtOH+THC: 0; EtOH+Vehicle: 3; Air+THC: 6; Air+Vehicle: 3).

### 3.2 Body Weights during Vapor Inhalation

Maternal body weights were recorded each morning **prior to** beginning the vapor inhalation paradigm. Since dams typically gave birth on GD 22, data from this day were excluded from all analyses (but shown in Figure 2 for reference). Dams exposed to vaporized EtOH gained significantly less weight over gestation (F[1,43] = 5.7, *p* < 0.05); however, prenatal THC exposure had no effect on the percentage of weight gain. To determine whether this difference was due specifically to vaporized EtOH exposure, body weight data were analyzed separately for the period prior to (GD 0-5) and following the initiation of vapor inhalation (GD 6-21). Note that since body weight recordings took place prior to vapor inhalation each day, body weight data from GD 5 were considered baseline and body weight on GD 21 (the morning after the last day of vapor exposure) were included in the analyses of body weights during drug exposure.

**Figure 2.**
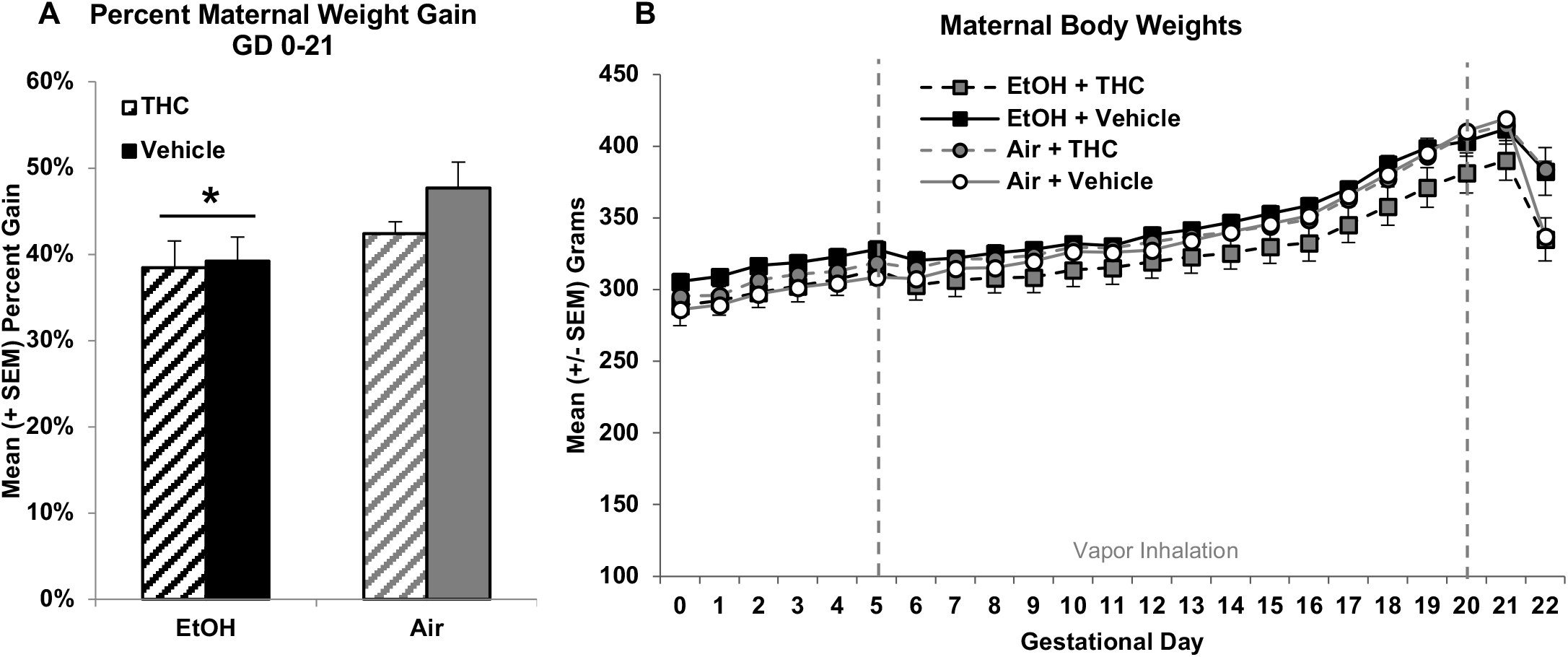
Although there were no statistically significant baseline differences in body weight, vaporized EtOH exposure slowed gestational weight gain, and THC did not alter this effect (A). THC exposure, by itself, did not significantly alter body growth (B). * = EtOH < Air, *p* < 0.05.

Baseline weights and weight gain over GD 0-5 (pre-vapor exposure) did not differ significantly among groups. However, during vapor exposure (GD 6-21), subjects exposed to EtOH grew at a slower rate, producing a main effect of EtOH (F[1,43] = 5.0, *p* < 0.05; Figure 2A) and a Day*EtOH interaction (F[15,145] = 3.4, *p* < 0.001; Figure 2B). There was also a significant interaction of Day*THC (F[15,145] = 2.3, *p* < 0.01). However, there were no significant differences in body growth between the THC alone group (Air+THC) and controls (Air+Vehicle), indicating that this interaction was driven by the lag in weight gain seen in the combination exposure group (EtOH+THC).

### 3.3 Food and Water Intake

To determine if drug exposure altered food or water intake, food and water levels for each day were recorded. Data were averaged separately for pre-vapor inhalation intake (GD 0-4) and post-vapor inhalation intake (GD 5-20, collapsed every 4 days: GD 5-8, GD 9-12, GD 13-16, GD 17-20), for simplicity of presentation.

Prior to vapor inhalation, pregnant dams assigned to the THC exposure groups ate more food on average (F[1,43] = 8.0, *p* < 0.05), an artifact of random assignment. However, this effect was not evident during the vapor inhalation period, as there were no significant differences in food intake among groups (Figure 3A). Similar to food consumption, there were some differences in water consumption at baseline, despite random assignment. Although the interaction of EtOH and THC was not statistically significant, subjects exposed to the combination of EtOH+THC drank more at baseline, producing main effects of EtOH (F[1,43] = 5.2, p < 0.05) and THC (F[1,43] = 19.0, *p* < 0.01). In contrast, during the vapor inhalation period, Vehicle controls (Air+Vehicle) drank less than all 3 drug-exposed groups, producing a 3-way interaction of Day*EtOH*THC (F[3,129] = 3.5, *p* < 0.05); there were no significant differences in water intake among the drug-exposed groups on any given Day (Figure 3B). Importantly, there were no significant group differences when analyzing differences scores between water intake at the end of treatment compared to the beginning (GD 0). Thus, it is not clear if the increased water intake was due to vapor exposure or simply due to initial variation introduced during random assignment. Regardless, the data suggest that our drug exposure paradigm did not lead to changes in food or water intake.

**Figure 3.**
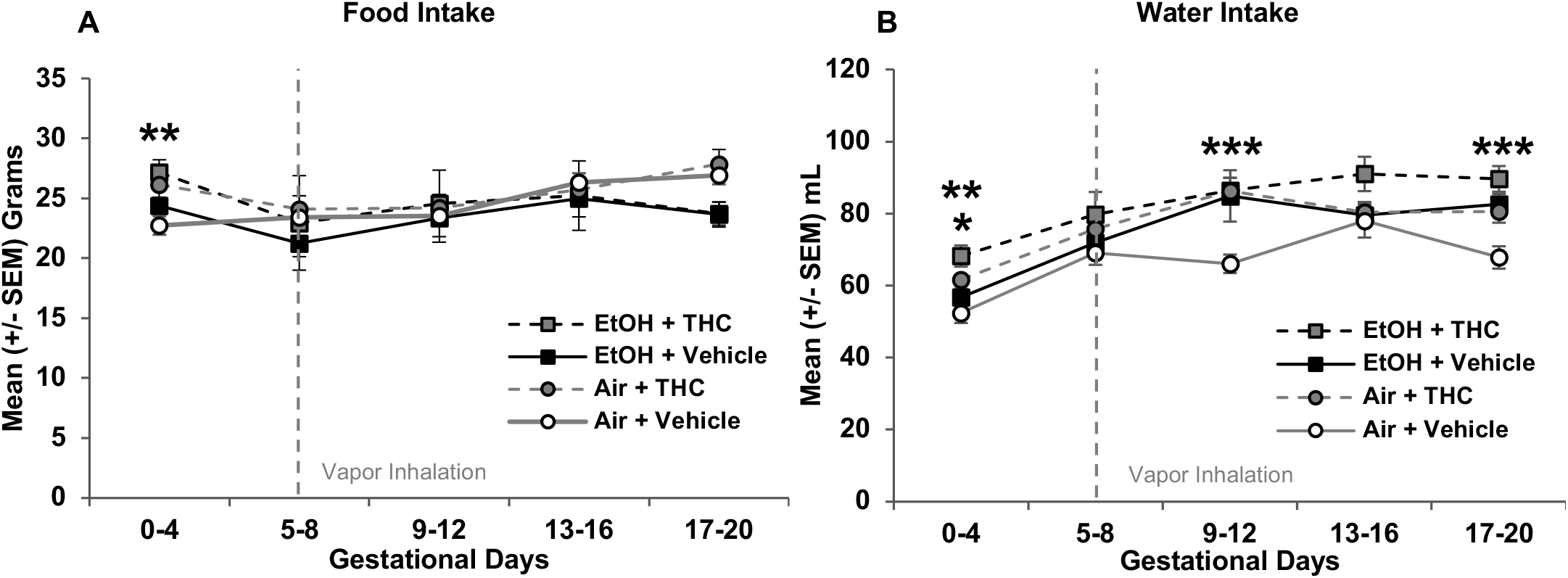
Exposure to EtOH or THC did not significantly alter food intake, although dams assigned to receive THC exposure ate more food than their counterparts at baseline (A). Dams exposed to EtOH or THC drank more than controls during vapor inhalation, although it is not clear if this was due to drug exposure or variation related to random group assignment (B). * = EtOH > Air (collapsed across THC group), *p* < 0.05. ** = EtOH+THC and Air+THC > Air+Vehicle, *p* < 0.05. *** = Air+Vehicle < all other groups, *p*’s < 0.01.

### 3.4 Core Body Temperatures

Previous studies have shown that THC exposure can reduce body temperature (Javadi-Paydar et al., 2018; Nguyen et al., 2016). In this study, only the group exposed to the combination of EtOH+THC exhibited significant reductions in body temperature during the exposure period (temperature change post-exposure minus pre-exposure), producing a main effect of THC (F[1,31] = 16.2, *p* < 0.001) and an interaction of EtOH*THC that neared significance (F[1,31] = 4.0, *p* = 0.055; Figure 4A). Interestingly, tolerance to this effect is seen over the course of exposure. Overall, dams exposed to the combination of EtOH+THC exhibited greater temperature reductions than all other groups (F[3,31] = 6.1, *p* < 0.01; SNK *p*’s < 0.05; Figure 4A). The 2-way interaction of Days*Group was near significance (F[9,93] = 1.8, *p* = 0.073). Follow-up analyses show that dams exposed to EtOH+THC had a greater temperature drop compared to all groups from GD 5-8 (F[3,31] = 4.2, *p* < 0.05; SNK *p* < 0.05) and 13-16 (F[3,31] = 7.5, *p* < 0.01; SNK *p* < 0.05), compared to EtOH+Vehicle and Air+Vehicle groups from GD 9-12 (F[3,31] = 5.3, *p* < 0.01; SNK *p* < 0.05), but only compared to EtOH+Vehicle dams during the final Days (GD 17-20; F[3,31] = 3.5, *p* < 0.05; SNK *p* < 0.05; Figure 4B).

**Figure 4.**
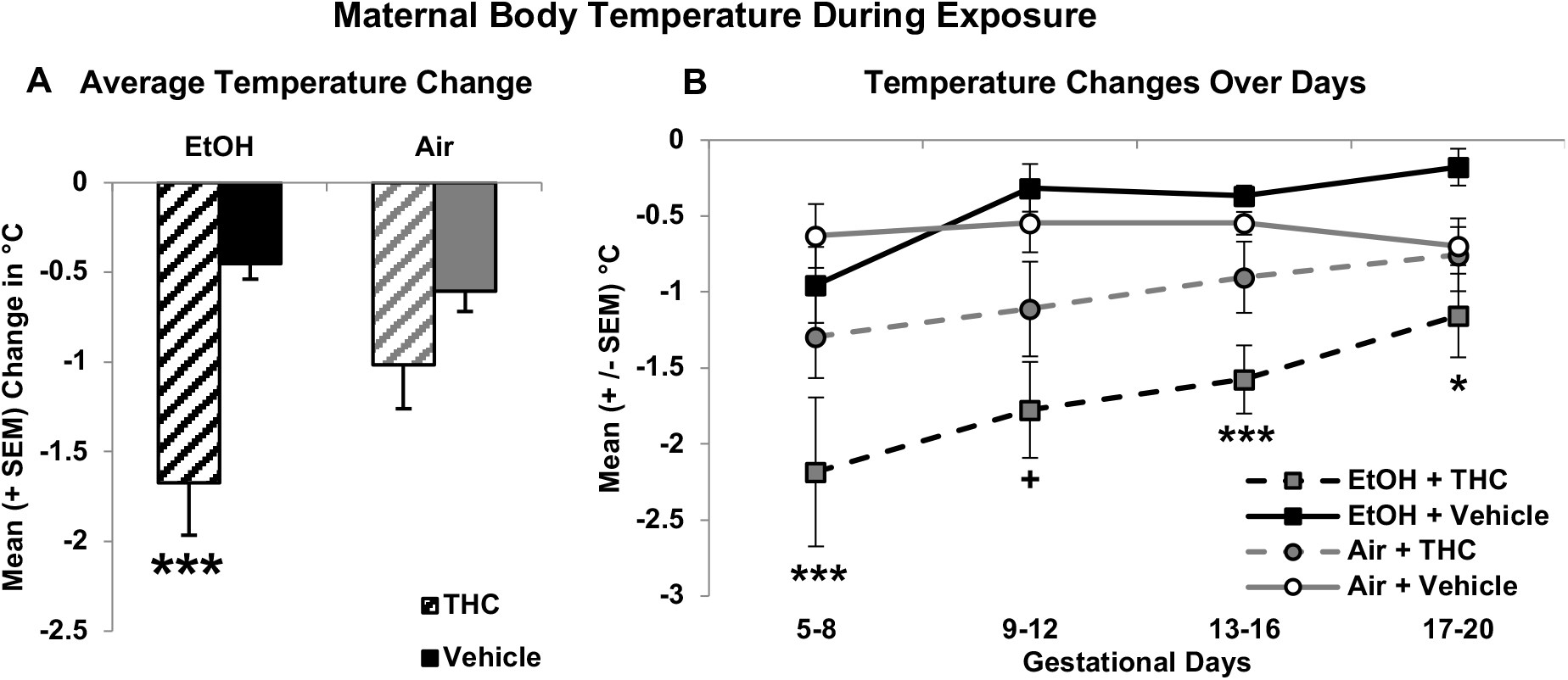
Vaporized THC exposure reduced body temperature during the exposure session when combined with ethanol (A). Interestingly, tolerance to this effect is observed over the course of exposure (B). *** = EtOH +THC < all other groups, *p*’s < 0.05. ^+^ = EtOH+THC < EtOH+Vehicle and Air+Vehicle, *p* < 0.05. * = EtOH+THC < EtOH+Vehicle, p < 0.05.

### 3.5 Blood Alcohol Concentrations

Blood samples for blood alcohol concentration (BAC) analyses were collected from all subjects, though only samples from the EtOH-exposed dams were analyzed. BACs (mg/dL) were initially analyzed using a 4 (day: GD5, GD10, GD15, GD20) × 5 (time: 15, 30, 60, 90, 180) × 2 (THC: EtOH+THC, EtOH+Vehicle) repeated measures ANOVA. Plasma samples from 5 subjects were not viable (EtOH+THC: 2, EtOH: 3). Overall, subjects exposed to the combination of EtOH+THC had higher BACs compared to those exposed to EtOH alone (EtOH+Vehicle; F[1,15] = 4.7, *p* < 0.05). As expected, BAC significantly declined across time (F[4,60] = 196.9, *p* < 0.001), but the decline was not altered by THC exposure. Rather, THC increased BAC at all time points: 15 (F[1,19] = 4.0, *p =* 0.06), 30 (F[1,19] = 4.6, *p <* 0.05), 60 (F[1,19] = 5.4, *p <* 0.05), 90 (F[1,19] = 4.6, *p <* 0.05), and 180 minutes (F[1,19] = 5.2, *p <* 0.05) post-vapor inhalation across days (Figure 5). A 2-way interaction of Day*Time was also evident (F[12,180] = 1.8, *p <* 0.05), due to slight variations across days. Data for each day and time are shown in Table 1.

**Table 1.**
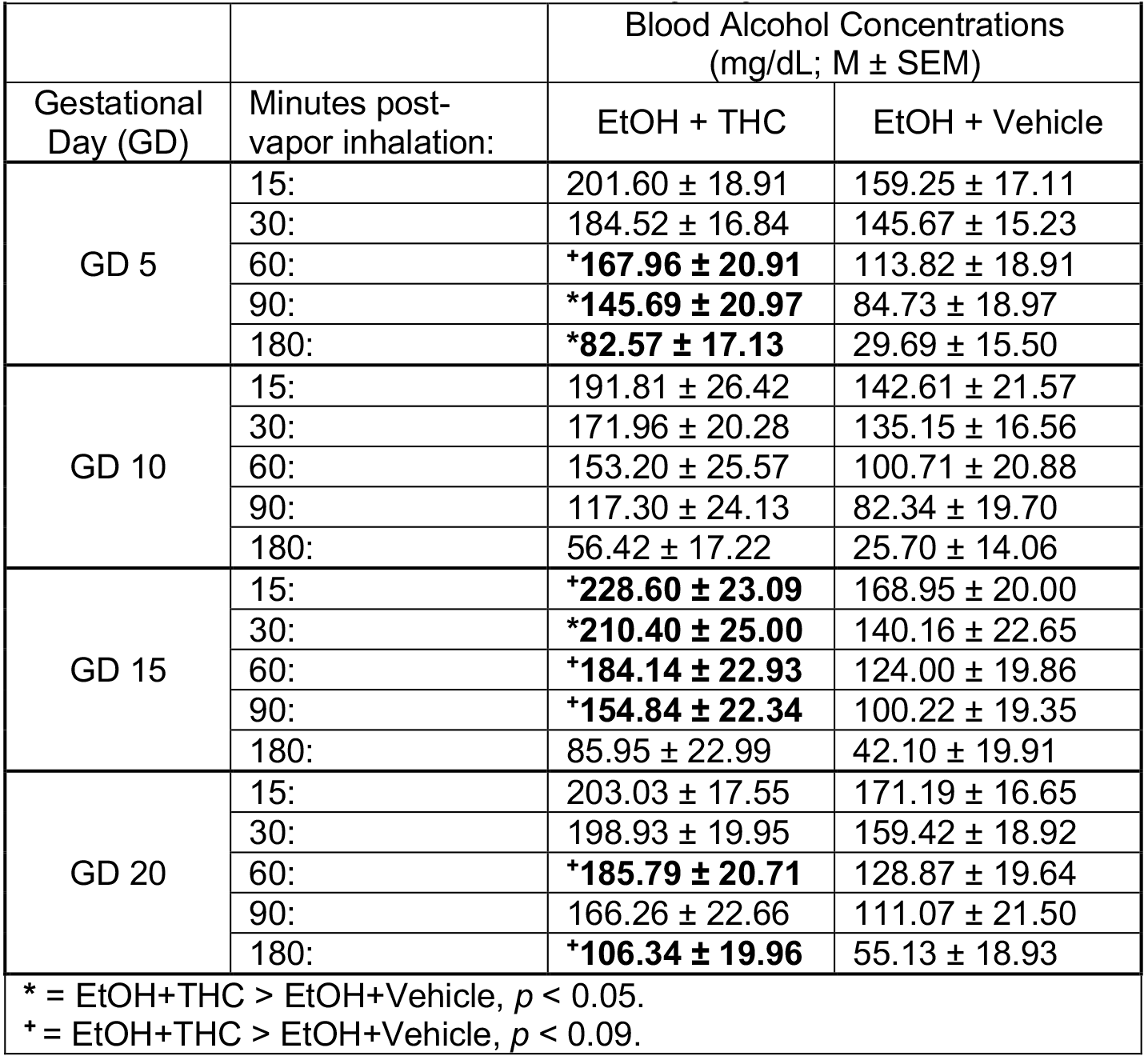
**Blood Alcohol Concentrations by Day and Time Point**. Dams exposed to the combination of EtOH+THC e-cigarette vapors had higher blood alcohol concentrations compared to those exposed to EtOH alone.

**Figure 5.**
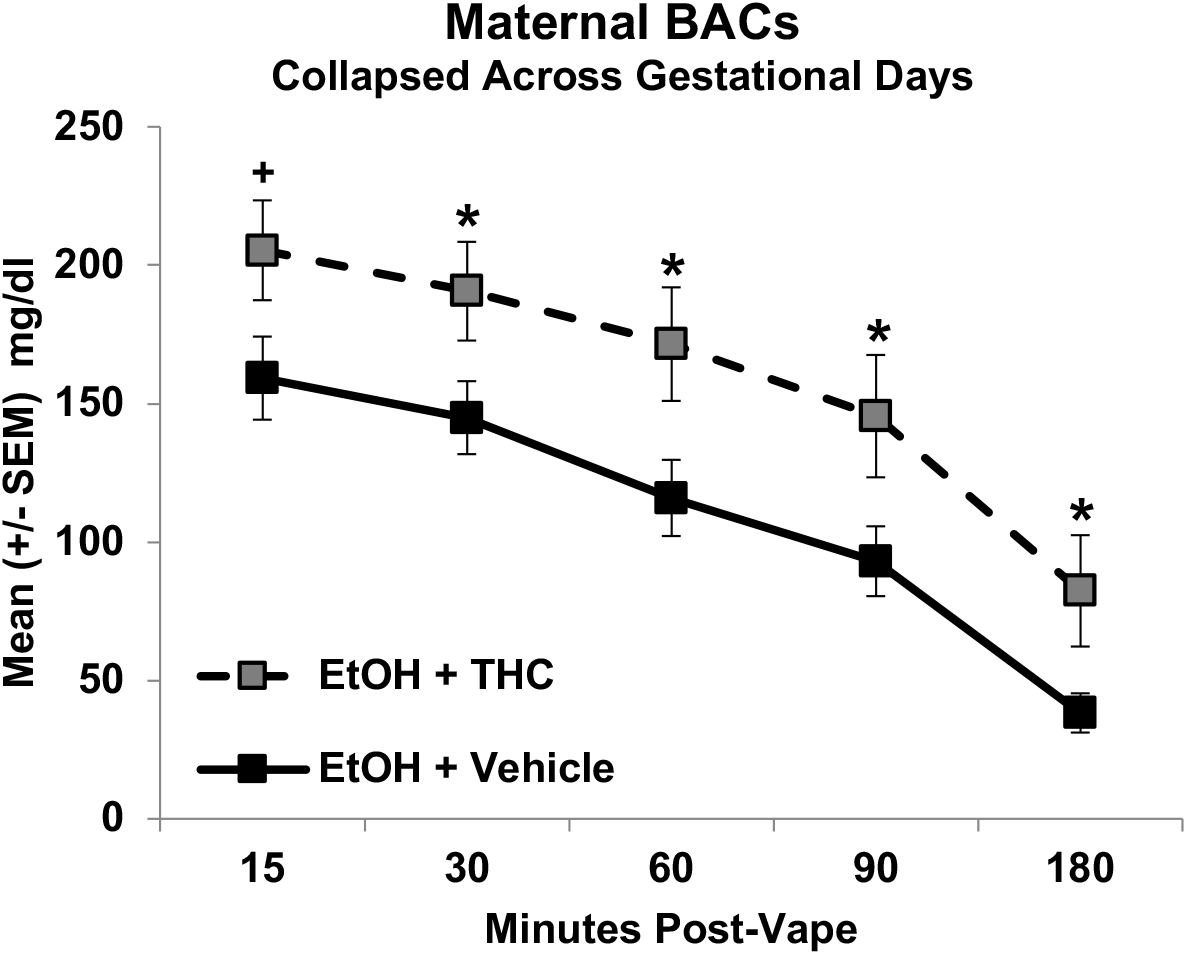
Combined exposure to vaporized ethanol and THC during gestation increased maternal blood alcohol concentrations more than exposure to ethanol alone at each time point. * = EtOH+THC > EtOH+Vehicle, *p*’s < 0.05. ^+^ = EtOH+THC > EtOH+Vehicle, *p* < 0.06.

### 3.6 THC and Metabolite Levels

Blood samples for THC and metabolite analyses (ng/mL) were collected from all subjects, though only samples from the THC-exposed dams were analyzed. Unfortunately, technical difficulties with our initial analyses reduced the number of viable samples from the EtOH+THC Group on GD 5 and 20; thus, only data from the THC alone group were analyzed for changes across Time on GD 5 and 20. Data from both groups were analyzed across time on GD 10 and 15; final data analyses included data from 13 dams (EtOH+THC: 6, THC: 7). THC and metabolite levels were analyzed using 5 (Time: 15, 30, 60, 90, 180) × 2 (EtOH: EtOH+THC, Air+THC [when possible]) repeated measures ANOVAs for each Day.

On the first day of vapor inhalation (GD 5), THC levels declined over the 180-minute time course (F[4,24] = 7.3, *p* < 0.05), as expected (Figure 6A). Metabolite levels also changed over time for both THC-OH (F[4,24] = 3.0, *p* < 0.05) and THC-COOH (F[4,24] = 35.1, *p* < 0.001; Table 2). Similarly, on GD 10, plasma THC levels declined over time (F[4,40] = 16.8, *p* < 0.001), and there was no significant difference between groups (Figure 6B). However, a significant interaction of Time*Group was observed in the THC-OH metabolite levels (F[4,44] = 4.1, *p* < 0.01). Although THC-OH levels declined over time for both groups (EtOH+THC: F[4,20] = 25.0, *p* < 0.001; Air+THC: F[4,24] = 10.4, *p* < 0.01), dams exposed to the combination of EtOH+THC had higher THC-OH levels than those exposed to THC alone, producing a main effect of Group (F[1,11] = 20.5, *p* < 0.01) and significant differences between groups at each time point (*p*’s < 0.05). No group differences were observed in THC-COOH levels, as similar patterns over time were observed (F[4,44] = 15.4, *p* < 0.001).

**Table 2.**
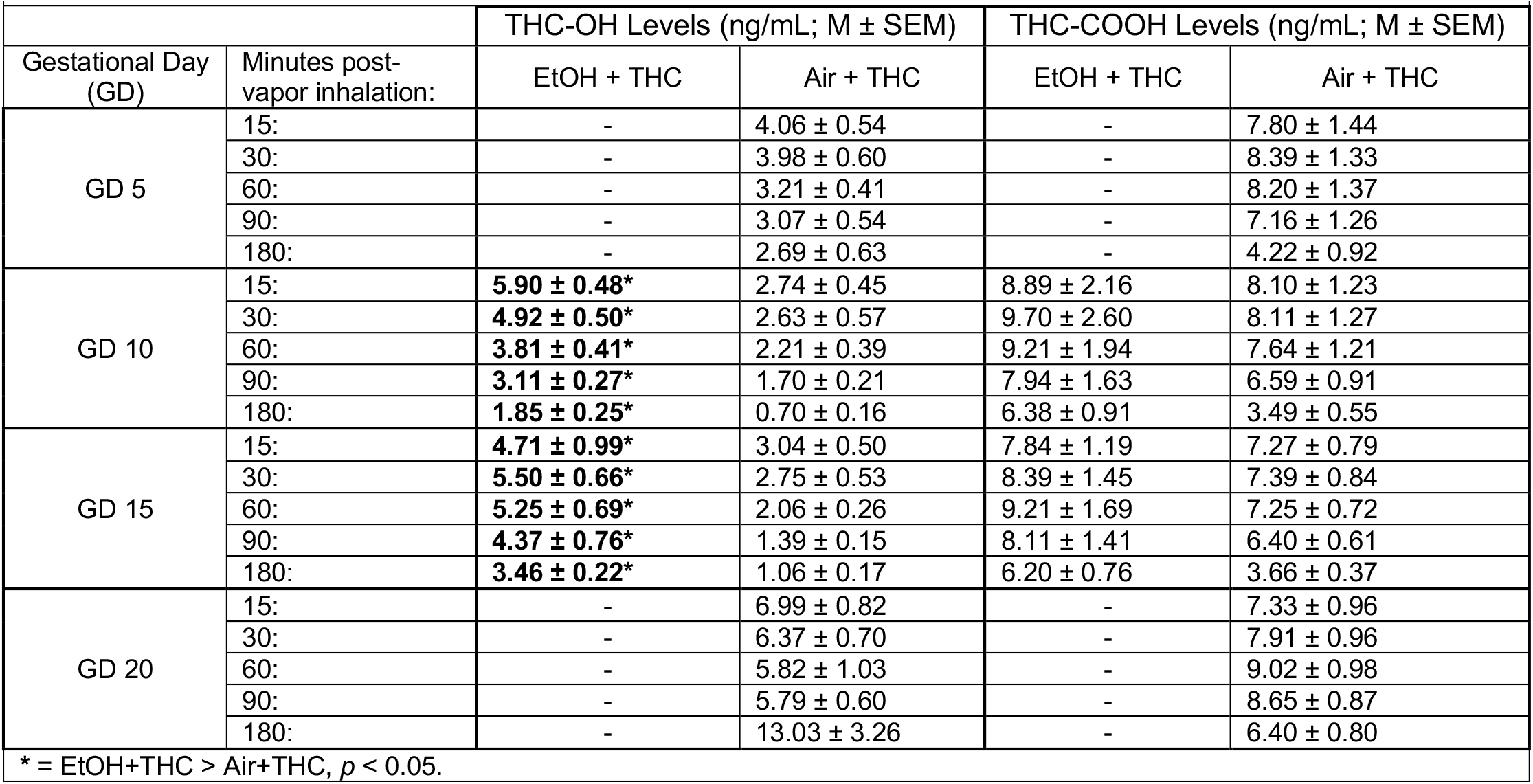
**THC Metabolite Levels by Day and Time Point**. Dams exposed to the combination of EtOH+THC e-cigarette vapors had higher THC-OH metabolite levels compared to those exposed to THC alone on gestational days 10 and 15.

| **Table 2. THC Metabolite Levels by Day and Time Point.** |  |  |  |  |  |
| --- | --- | --- | --- | --- | --- |
|  |  | THC-OH Levels (ng/mL; M ± SEM) |  | THC-COOH Levels (ng/mL; M ± SEM) |  |
| Gestational Day (GD) | Minutes post-vapor inhalation: | EtOH + THC | Air + THC | EtOH + THC | Air + THC |
| GD 5 | 15: | - | 4.06 ± 0.54 | - | 7.80 ± 1.44 |
|  | 30: | - | 3.98 ± 0.60 | - | 8.39 ± 1.33 |
|  | 60: | - | 3.21 ± 0.41 | - | 8.20 ± 1.37 |
|  | 90: | - | 3.07 ± 0.54 | - | 7.16 ± 1.26 |
|  | 180: | - | 2.69 ± 0.63 | - | 4.22 ± 0.92 |
| GD 10 | 15: | **5.90 ± 0.48\*** | 2.74 ± 0.45 | 8.89 ± 2.16 | 8.10 ± 1.23 |
|  | 30: | **4.92 ± 0.50\*** | 2.63 ± 0.57 | 9.70 ± 2.60 | 8.11 ± 1.27 |
|  | 60: | **3.81 ± 0.41\*** | 2.21 ± 0.39 | 9.21 ± 1.94 | 7.64 ± 1.21 |
|  | 90: | **3.11 ± 0.27\*** | 1.70 ± 0.21 | 7.94 ± 1.63 | 6.59 ± 0.91 |
|  | 180: | **1.85 ± 0.25\*** | 0.70 ± 0.16 | 6.38 ± 0.91 | 3.49 ± 0.55 |
| GD 15 | 15: | **4.71 ± 0.99\*** | 3.04 ± 0.50 | 7.84 ± 1.19 | 7.27 ± 0.79 |
|  | 30: | **5.50 ± 0.66\*** | 2.75 ± 0.53 | 8.39 ± 1.45 | 7.39 ± 0.84 |
|  | 60: | **5.25 ± 0.69\*** | 2.06 ± 0.26 | 9.21 ± 1.69 | 7.25 ± 0.72 |
|  | 90: | **4.37 ± 0.76\*** | 1.39 ± 0.15 | 8.11 ± 1.41 | 6.40 ± 0.61 |
|  | 180: | **3.46 ± 0.22\*** | 1.06 ± 0.17 | 6.20 ± 0.76 | 3.66 ± 0.37 |
| GD 20 | 15: | - | 6.99 ± 0.82 | - | 7.33 ± 0.96 |
|  | 30: | - | 6.37 ± 0.70 | - | 7.91 ± 0.96 |
|  | 60: | - | 5.82 ± 1.03 | - | 9.02 ± 0.98 |
|  | 90: | - | 5.79 ± 0.60 | - | 8.65 ± 0.87 |
|  | 180: | - | 13.03 ± 3.26 | - | 6.40 ± 0.80 |
| * = EtOH+THC > Air+THC, *p* < 0.05. |  |  |  |  |  |

**Figure 6.**
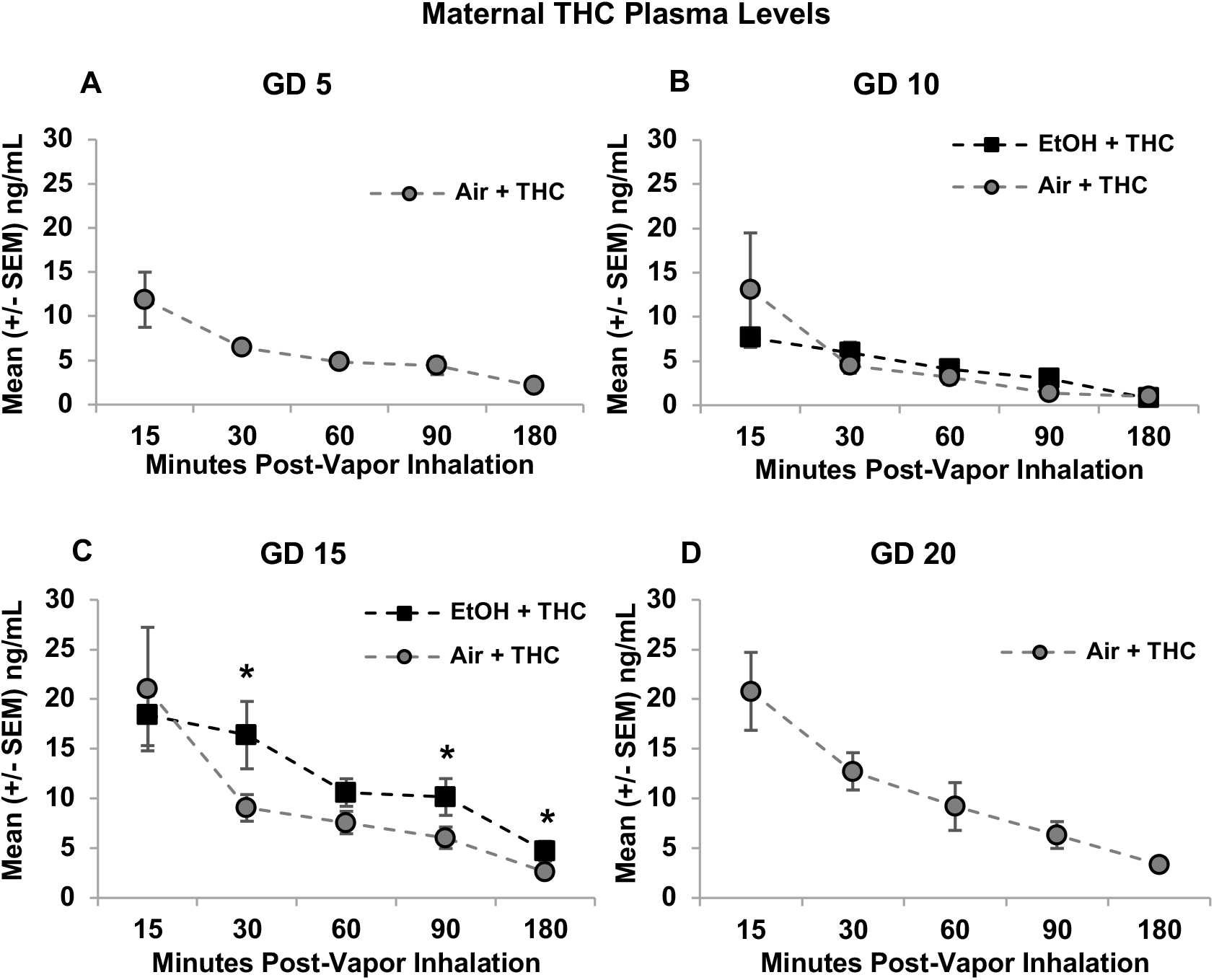
THC plasma levels declined over 180 minutes post-vapor inhalation, on each day of the vapor exposure (GD 5 [A], GD 10 [B], GD 15 [C], GD 20 [D]). Levels also gradually increased across days in both exposure groups. On GD 15, subjects exposed to the combination of EtOH and THC had higher plasma THC levels than those exposed to THC alone. * = EtOH+THC > Air+THC, *p*’s < 0.05.

By GD 15, however, dams exposed to the combination of EtOH+THC had higher THC plasma levels than dams exposed to Air+THC, specifically at 30 (F[1,11] = 6.5, *p* < 0.05), 90 (F[1,11] = 6.8, *p* < 0.05), and 180 minutes post-vape (F[1,11] = 10.3, *p* < 0.01; Figure 6C). Overall, THC levels declined over Time (F[4,36] = 4.4, *p* < 0.01). Similarly, THC-OH metabolite levels declined over time (F[4,40] = 22.6, *p* < 0.001), but pregnant dams exposed to the combination of EtOH+THC had higher THC-OH levels overall (F[1,10] = 14.3, *p* < 0.01) and at each time point (*p*’s < 0.05). THC-COOH levels also declined (F[1,10] = 36.9, *p* < 0.001), but no group differences were observed.

On the last Day of vapor inhalation (GD 20), levels of plasma THC (F[4,24] = 24.4, *p* < 0.001), THC-OH (F[4,24] = 3.4, *p* < 0.05), and THC-COOH (F[4,24] = 18.0, *p* < 0.001) changed over time in dams exposed to Air+THC (Figure 6D), as expected. Importantly, plasma THC levels at 15 minutes post-inhalation increased from GD 5 to 20 (F[1,6] = 6.3, *p* < 0.05), from ∼11 to over 20 ng/mL. In fact, even though data for dams exposed to the combination of EtOH+THC were unavailable for GD 5 and 20, a significant increase in plasma THC levels was observed from just GD 10 to 15 within this group (F[1,5] = 10.9, *p* < 0.05). However, neither THC-OH nor THC-COOH metabolite levels showed similar increases over days in either group.

### 3.7 Gestational Length and Litter Composition

Gestational length was not significantly affected by prenatal EtOH, THC, or combined exposure (data not shown). In addition, no differences were observed in the number of total pups born, or the male:female ratio of the litter. Lastly, the average pup weight on PD 2 (day of culling) was not significantly altered by prenatal EtOH, THC, or combined exposure (Table 3).

**Table 3.** **Litter Composition**. Means (±SEM) of litter composition by prenatal exposure group. Prenatal exposure to EtOH, THC, or the combination did not alter the number of offspring, the male:female ratio, or offspring weights compared to the controls.

| **Table 3. Litter Composition.** |  |  |  |
| --- | --- | --- | --- |
| Litter Exposure | Number of Offspring<br>M ± SEM | Male:Female Ratio<br>M ± SEM | Weight (g)<br>M ± SEM |
| EtOH + THC (n=10) | 10.70 ± 0.99 | 0.96 ± 0.23 | 7.46 ± 0.24 |
| EtOH + Vehicle (n=12) | 12.08 ± 0.91 | 0.92 ± 0.21 | 7.65 ± 0.27 |
| Air + THC (n=13) | 12.23 ± 0.87 | 1.15 ± 0.20 | 7.69 ± 0.26 |
| Air + Vehicle (n=13) | 12.25 ± 0.91 | 1.27 ± 0.21 | 8.19 ± 0.27 |

### 3.8 Offspring Development

#### 3.8.1 Eye Opening

Shapiro-Wilk normality tests indicated that the eye opening data for the offspring were not normally distributed (*p*’s < 0.05) due to the limited numerical range of outcomes. Thus, non-parametric analyses were used. Females and males did not significantly differ on the first day of eye opening, so non-parametric analyses of eye opening included both male and female offspring.

Both a Mann-Whitney U (EtOH, Air) and Kruskal-Wallis H (Exposure Group) analyses indicated that prenatal EtOH exposure via vapor inhalation significantly delayed the first day of eye opening (U = 607.7, *p* < 0.01; H = 12.4, *p* < 0.01; Figure 7). Prenatal THC exposure did not significantly alter this developmental milestone (U = 923.5, *p* = 0.70).

**Figure 7.**
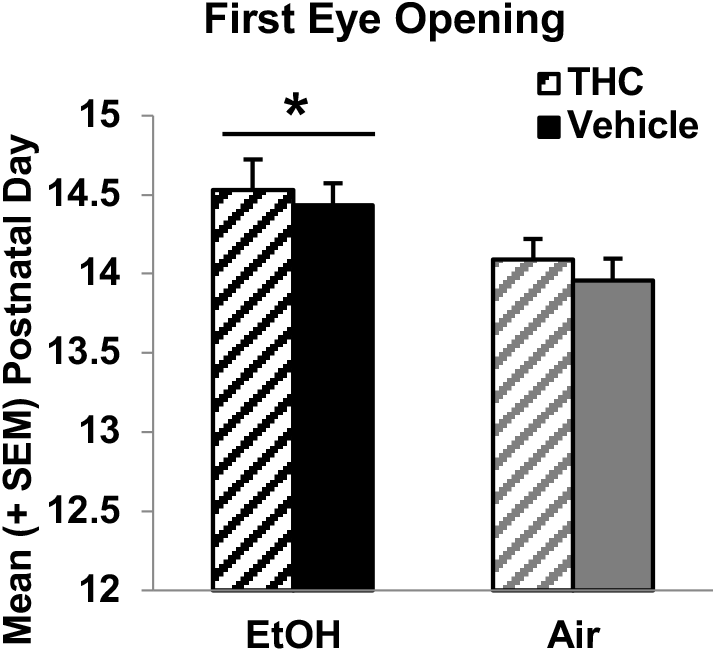
Prenatal EtOH exposure delayed eye opening. * = EtOH > Air, *p* < 0.01.

#### 3.8.2 Adolescent Body Weights

By PD 30, offspring exposed to prenatal THC weighed less than Vehicle-exposed groups (F[1,79] = 6.3, *p* < 0.05; Figure 8). Although there were no significant interactions between either drug exposure or sex, this THC-related effect was slightly more robust in the male offspring (F[1,38] = 4.0, *p* = 0.05). As expected, females weighed less than males (F1,79] = 30.3, *p* < 0.001). Offspring exposed to EtOH did not differ significantly in weight from Air-exposed groups.

**Figure 8.**
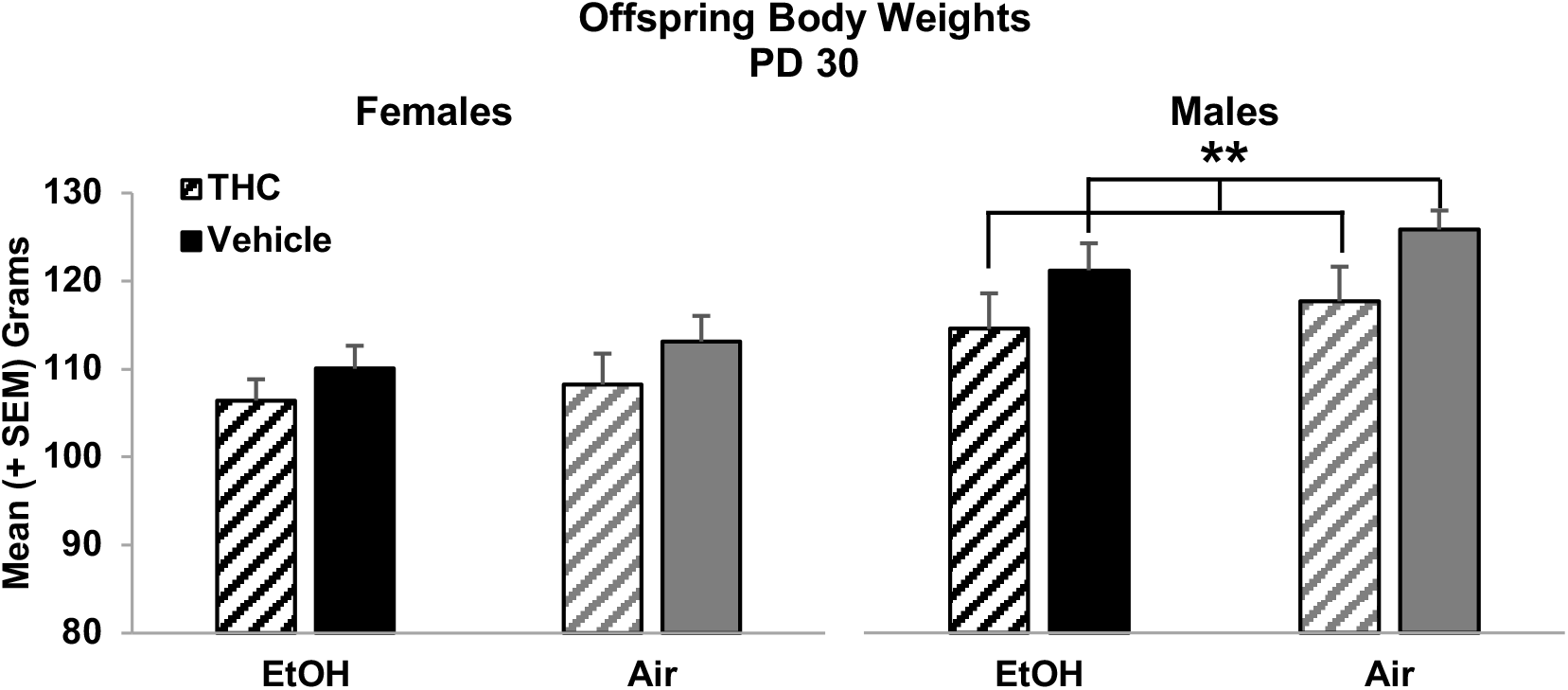
By adolescence, offspring exposed to THC e-cig vapors prenatally weighed significantly less than Vehicle-exposed offspring. Although there was no interaction with sex, this effect was more robust among male offspring. ** = THC < Vehicle, *p* = 0.05.

## 4 Discussion

To our knowledge, this is the first study to use a clinically relevant e-cigarette model of THC exposure in combination with ethanol, mimicking a common polydrug use seen among pregnant women. Moreover, this is the first study to closely monitor multiple maternal factors and early offspring measures following cannabis e-cigarette exposure. Importantly, this study demonstrated that co-exposure to THC elevates blood alcohol levels during pregnancy. In addition, THC levels rise over the course of exposure during the pregnancy and THC metabolism is altered by alcohol. These pharmacokinetic interactions may have important implications for fetal risk to combined exposure.

The timing and administration of THC in this study models common use patterns among pregnant women. We exposed pregnant Sprague-Dawley rats to alcohol and/or THC e-cigarette vapors daily from GD 5-20, a period equivalent to the first and second trimesters in humans. Cannabis consumption during pregnancy is especially prevalent during the first and second trimesters (Volkow et al., 2019). During early pregnancy, cannabis is often used to combat undesirable symptoms such as nausea, or used as an alternative to other medications under the belief that cannabis use during pregnancy is safer (Brown et al., 2017). Moreover, our paradigm administered THC to pregnant rats using commercially available e-cigarettes, which are a popular route of administration for cannabis products.

Importantly, we found an interactive effect of alcohol and THC on blood alcohol concentrations. Blood alcohol concentrations in dams exposed to alcohol peaked around 150 mg/dL, whereas pregnant dams exposed to the combination of alcohol and THC had significantly higher blood alcohol concentrations, peaking at 200 mg/dL. These levels translate to a moderate-high binge episode of alcohol consumption, roughly twice the legal limit of BAC for driving across most of the United States (0.08%; 80 mg/dL). One clinical study examining blood levels following co-use of alcohol and cannabis in adult users suggested that THC may lower blood alcohol concentrations (Lukas et al., 1992); however, in that study, THC was administered before alcohol (opposite of the current study), and the results have been challenged (Perez-Reyes et al., 1993). Most clinical adult research examining blood alcohol concentrations following co-use measure BAC prior to THC administration to focus on changes in THC levels; thus, this interaction has not been shown or examined before. Other research from our laboratory has shown a similar elevation in blood alcohol level with combined alcohol and a synthetic cannabinoid (CP-55,940); however, this previous study used a 3^rd^ trimester exposure model, and thus subjects were neonates rather than pregnant dams (Breit et al., 2019b). In any case, preclinical data have consistently shown that higher exposure to alcohol during early development leads to more extreme consequences (Burd et al., 2012), and thus this is an important synergistic effect to consider.

Relatedly, THC levels increased with repeated exposure over days, increasing from 10-15 ng/mL to over 20 ng/mL during the course of pregnancy. Thus, THC appears to accumulate in the dams throughout the vapor inhalation exposure period, as levels rose in both groups over days. This is a novel finding in teratology research, but has been shown repeatedly in models of chronic cannabis exposure in adult rats (Fleischman et al., 1975). This suggests that THC may not be completely excreted by pregnant rodents over a 24-hour period; thus, repeated exposure may lead to increased THC plasma levels as days progress, and may not produce comparable plasma levels to acute or intermittent exposure models. Moreover, the combination of alcohol altered THC metabolism.

Unfortunately, we did not have adequate THC data from GD 5 and 20 in the combined exposure group. Nevertheless, we saw that pregnant dams exposed to the combination of alcohol and THC had similar THC levels than those exposed to THC alone on GD 10, but significantly higher THC levels on GD 15. Dams exposed to the combination of alcohol and THC also had higher THC-OH metabolite levels on both GD 10 and 15. These data suggest that co-exposure with alcohol alters THC metabolism. THC levels from the combination of subjects exposed to both EtOH and THC on GD 20 would illuminate whether the trajectory of combined effects persisted to the end of the pregnancy. Either way, it will be critical for future studies to understand the pharmacokinetic interactions of these drugs and the implications for fetal development.

It is unclear exactly why these interactions in blood levels of alcohol and THC are present. One possibility is that co-exposure via vapor inhalation altered overall metabolism in pregnant subjects. A recent study showed that adolescent male rats exposed to alcohol (via drinking) and THC (via s.c. injection) exhibited altered glucose and insulin homeostasis, which could affect metabolism rates (Nelson et al., 2019); however, there are many obvious methodological differences between these two study designs, and this previous work did not yield differences in blood alcohol levels. Given the reduced body temperatures during sessions among the combined exposure group, it would be possible that lower temperature could influence overall drug metabolism. However, this also seems unlikely, given that the body temperature reductions declined across days, a pattern neither consistent with elevations in BAC nor THC.

The most probable possibility is that alcohol exposure increased absorption of THC; when alcohol precedes cannabis exposure, alcohol absorption can increase plasma THC levels in humans (Lukas et al., 2001) and prolong elimination (Toennes et al., 2011). These interactions are consistent with what has been documented in adult users in clinical studies, where alcohol can increases THC when alcohol is consumed first (Downey et al., 2013; Hartman et al., 2015). Interestingly, we found that EtOH also elevates THC-OH whether THC levels are elevated (GD 15) or not (GD 10); THC-OH is the main psychoactive metabolite of THC. Indeed, clinical research in adult users has also shown increased THC-OH levels following alcohol consumption (Hartman et al., 2015). In contrast, THC-COOH is the second main metabolite of THC and is non-psychoactive. Ethanol did not significantly alter THC-COOH levels, perhaps because our THC plasma levels were moderate, because the time course of THC-COOH elevations are delayed and group differences may only be seen at later time points, or because of differences in elimination routes of THC-OH vs. THC-COOH. Nevertheless, elucidation of how alcohol modifies THC absorption and metabolism is critical for understanding the consequences of polydrug administration.

In order to obtain immediate confirmation that THC was inducing a physiological effect, we recorded core body temperatures of each dam before and after vapor inhalation each day. Previous work shows that THC vapor decreases core temperatures in female, non-pregnant rats (Javadi-Paydar et al., 2018; Nguyen et al., 2016). Although we did not find significant reductions in body temperature among dams exposed only to THC e-cigarette vapor exposure, there were significant temperature reductions in dams exposed to both THC and alcohol, another synergistic effect. Interestingly, the body temperature drops became less robust over the course of exposure, indicating tolerance, despite increasing THC levels.

Unfortunately, prenatal, preclinical studies rarely report THC levels, so it is challenging to compare our levels to other studies, particularly since this is a novel administration route. Our THC levels represent a low-moderate exposure, similar to studies used in human adults (Hartman et al., 2015). We should point out that blood sampling began at 15 minutes after removal from the chamber, and following a 10-minute air clearance period. Thus, it is possible that THC levels were higher prior to our first sampling period. Nevertheless, given that blood THC levels of humans reach around 100 ng/mL following consumption of marijuana cigarettes with moderate-high levels of THC (Andrenyak et al., 2017), our levels are physiological and clinically relevant to occasional use of low-moderate THC level products among pregnant women. It should be emphasized that there are a wide variety of cannabis products sold today, with both low and high potency levels of THC; thus, it is important that we research all levels of this constituent and not target toward only extreme levels.

Despite the pharmacokinetic interactions, only the alcohol-exposed dams significantly lagged in weight gain. The alcohol dose used in this study mimics a moderate binge (150 mg/dL), and although the current study used a lower dose and different administration route than past research in our lab, the effects on maternal weight gain are consistent (Thomas et al., 2009). In contrast, THC e-cigarette vapor did not alter maternal weight gain on its own, but body weights of dams exposed to the combination of alcohol and THC did lag, though statistical analyses were not significant. Importantly, this effect was not due to altered food intake. Nevertheless, food intake should be recorded in studies of prenatal drug exposure and it will be important to determine whether pair-fed controls would be necessary at higher doses of either drug.

Despite the effects observed in body weight gain, blood alcohol levels, and THC levels, we did not observe any significant changes in the pregnancy health, gestational length, sex ratio or birth weight. Previous clinical and preclinical research has shown that prenatal alcohol exposure can alter several pregnancy outcome variables (Bailey et al., 2011), but limited clinical data suggests that prenatal cannabis exposure does not significantly increase the likelihood of pre-term delivery or miscarriages (Fried et al., 2001; Gunn et al., 2016; Huizink, 2014). In the current study, we did not observe a higher rate of miscarriage, dystocia, or cannibalization in any group; in fact, there were zero instances of complications in the combination group (EtOH+THC). This finding contrasts with previous data from our lab reporting higher mortality rates in neonates exposed to alcohol and CP-55,940, although this previous study used much higher doses of each drug and directly intoxicated the neonate rather than the dam (Breit et al., 2019b). Importantly, although this co-exposure model using vapor inhalation does not directly affect maternal and litter viability, it does produce long-term behavioral effects in development among offspring (Breit et al., 2019a, 2020), which will be reported separately.

Very recently published data using preclinical models show that prenatal exposure to THC leads to severe fetal growth restrictions (Natale et al., 2020), although that study utilized an i.p. injection route. Although we did not observe changes in litter size or average pup weight on PD 2 using this administration route and doses of alcohol and THC, we did find that prenatal THC exposure reduced body weights in offspring at PD 30. Thus, it is possible that gross differences in weight may not be observed until later in life. It is also possible that prenatal exposure to THC may have altered puberty onset which could have attributed to weight differences on PD 30; puberty onset was not measured in the current study. In addition, we recorded the first day of eye opening, which is a developmental milestone in rodents. While we did not find that prenatal THC e-cig exposure altered the first day of eye opening, we did observe that prenatal alcohol exposure delayed eye opening, consistent with past literature (Thomas et al., 2009).

There are several limitations in the current study to address. First, only single doses of each drug were used to examine initial effects and establish this co-consumption vapor inhalation model. Although the doses of each drug were low to moderate, we did find interactive effects even at these levels that are commonly consumed, which is an important finding with clinical significance. However, future research in this area should examine multiple doses in order to further elucidate dose-dependent effects that characterize the range of human consumption to better understand interactive effects of prenatal alcohol and THC exposure.

Furthermore, although THC is the primary psychoactive component of cannabis, it is by no means the only one. The cannabis plant contains more than 500 chemical compounds, including over 100 naturally-occurring cannabinoids (Radwan et al., 2017). For example, cannabidiol (CBD) has both cannabinoid-and non-cannabinoid-receptor mechanisms of action, and can elicit neurogenesis and neuroprotection instead of psychoactive effects on the user (Boggs et al., 2018; Madras, 2019). Often, cannabis products include a combination of THC and CBD; CBD can mitigate the psychological and cognitive effects of THC to make them more tolerable, although the mechanism by which this occurs is poorly understood (Boggs et al., 2018; Madras, 2019). We acknowledge that the outcomes presented in the current study may vary based on cannabinoid constituents.

It is also important to mention that while THC e-cigarette vapor exposure mimics current consumption patterns, vaporized alcohol exposure is not as clinically relevant. We chose that administrative route to minimize additional stress, since subjects would already be exposed to e-cigarette vapor. Our lab has extensively studied the effects of prenatal alcohol exposure using intragastric intubation at a higher dose of alcohol, and we have seen similar effects on maternal weight gain and eye opening (Thomas et al., 2009). Others have also found that vaporized alcohol yields similar consequences to traditional gastric administration routes in neonatal mice (Ryabinin et al., 1995). Importantly, the BAC achieved is the critical factor, particularly since alcohol and THC were not administered simultaneously. Moreover, we have previously reported synergistic effects of alcohol and cannabinoids (both on BAC and offspring behavior) when they are administered via different routes during the neonatal period (Breit et al., 2019b, 2019c), suggesting that synergistic effects are not reliant on a single route of administration. Thus, although this is the first study to administer polydrug exposure via vapor inhalation during the prenatal period, these factors collectively suggest that consequences of prenatal alcohol, as well as the synergistic effects of alcohol and THC, observed in the current study are not due to the route of administration.

We also want to acknowledge that the control group in this study was exposed to the e-cigarette vehicle, propylene glycol. Although not presented here, we did compare the data in this control group (Air+Vehicle) to a small pilot of non-handled, no-vape controls. We did not find any differences between these groups in maternal body weight, food or water intake, nor were there differences in gestation length, or any of the litter outcome variables. Nevertheless, little is known about whether vaping the vehicle constituents, by themselves, may exert damaging effects on physical and behavioral development of the fetus (Strongin, 2019).

In summary, these results suggest that this novel co-exposure vapor inhalation paradigm effectively induces physiological changes in pregnant dams and long-term alterations among offspring while avoiding nutritional confounds, birth complications, general neonate health, or maternal behaviors toward the offspring. With this model, we have found important pharmacokinetic interactions between alcohol and THC. We are currently examining the long-term effects of combined prenatal exposure to alcohol and cannabis constituents on long-term brain and behavioural development in the offspring. Given the high rate of co-use of these drugs during pregnancy, and that many births in the United States are unplanned (Mosher et al., 2012), understanding the effects of prenatal exposure to both of these drugs (individually and in combination) has important implications, not only for the lives of affected individuals and families, but also for public health and establishing public policy.

## 5 Acknowledgements

Supported by National Institute on Alcohol Abuse and Alcoholism grant AA025425 to Dr. Thomas and an NIH Loan Repayment Program award to Dr. Breit. Special thanks to Maury Cole at La Jolla Alcohol Research, Inc. (San Diego, CA) for building the vapor inhalation equipment and for co-exposure advisement. We also want to recognize Dr. Michael Taffe for his aid and advisement in the implementation of this vapor inhalation paradigm, as well as to Dr. Jacques Nguyen, Sophia Vandewater, and Kevin Creehan for training Dr. Breit on intravenous catheter implantation and blood collection procedures. Lastly, we want to recognize the member of the Center for Behavioral Teratology at San Diego State University for assisting in data collection and interpretation, particularly the instrumental efforts of Brandonn Zamudio, Samirah Hussain, and Ivette Gonzalez.

